# Directed-evolution mutations of adenine base editor ABE8e improve its DNA-binding affinity and protein stability

**DOI:** 10.1101/2023.05.09.539947

**Authors:** Haixia Zhu, Xinyi Jiang, Lei Wang, Qin Qin, Menghua Song, Qiang Huang

## Abstract

Adenine base editors (ABEs), consisting of CRISPR-associated (Cas) nickase and deaminase, can convert the A:T base pair to G:C. Previous studies have shown that ABE8e, a directed-evolution variant of ABE7.10 with eight amino-acid mutations in its deaminase TadA-8e, has higher base editing activity than ABE7.10. However, it remains unclear how the directed-evolution mutations of ABE8e increase its editing activity. Here, we combined molecular dynamics (MD) simulations and experimental measurements to elucidate the molecular origin of the activity enhancement by these mutations. MD simulations and microscale thermophoresis measurements showed that TadA-8e has a higher DNA-binding affinity than TadA-7.10, and the main driving force is electrostatic interactions. The directed-evolution mutations increase the positive charge density in the DNA-binding region, thereby enhancing the electrostatic attractions with DNA. We showed that R111 is the key mutation for the enhanced binding to DNA, which explains why the ABE-8e activity was dramatically reduced when R111 was mutated back to the original T. Unexpectedly, we also found that these mutations improve the thermal stability of TadA-8e by ∼12°C (T_m_). Our results indicate that the editing activities of ABEs are closely related to their DNA-binding affinity and protein stability, thus providing a rational basis for their optimization.

**Author Summary:** Adenine-to-guanine mutations account for 47% of known disease-causing point mutations in humans. Adenine base editors (ABEs), which can restore the guanine mutation to adenine, are a promising tool for precision gene therapy. ABE8e is the most widely used editor today due to its high editing efficacy and was derived from the first-generation base editor ABE7.10 by directed evolution. Compared to ABE7.10, ABE8e contains 8 directed-evolution amino-acid mutations. Understanding how these mutations affect the efficiency of ABE8e is important for the development of efficient base editors. In this study, we combined computational and experimental approaches to investigate how these mutations affect the editing activity of ABE8e. Our results showed that these directed-evolution mutations improve the DNA binding affinity and the protein stability of ABE8e, by enhancing electrostatic and hydrogen bonding interactions. Therefore, our study provides valuable insights for the design of more efficient base editors.

## Introduction

CRISPR systems have been widely used as molecular machines for targeted genome editing [1-4]. In general, Cas nucleases induce double-stranded breaks (DSBs) at target DNA sites, which are repaired by non-homologous end joining (NHEJ) and homology-directed repair (HDR), ultimately leading to genome modification [2, 5, 6]. However, NHEJ is an uncontrollable pathway that usually results in random insertion or deletion mutations (indels) [7, 8]. And the addition of a donor DNA template can stimulate HDR to precisely modify the gene, but this pathway occurs with low efficiency [9, 10]. Thus, although the CRISPR system is an efficient tool for disrupting genes, applications that modify bases at specific DNA sites require more precise DNA editing tools [11-13]. For example, the largest class of known human pathogenic mutations are point mutations, the correction of which requires very precise site-specific editing [14, 15].

To avoid the unwanted mutations, DNA base editors (BEs) have been developed to allow programmable conversion of the target base pairs without creating DSBs [16-18]. BEs consist of a Cas nickase (e.g., Cas9n) and a deaminase that acts on single-stranded DNA (ssDNA). After the single guide RNA (sgRNA) is paired with the target DNA strand (TS), a segment of the non-target DNA strand (NTS) becomes unpaired, and the DNA bases within this ssDNA segment are modified by the deaminase; at the same time, the Cas nickase cleaves the unedited TS, inducing cells to use the edited NTS as a template to repair the TS, and ultimately accomplishing the base editing (Fig. 1A)[16, 17, 19]. To date, two types of BEs have been discovered: cytosine base editors (CBEs), which convert a C:G base pair into T:A [16, 20], and adenine base editors (ABEs), which convert an A:T base pair to G:C [17, 21]. It is estimated that about 60% of human disease-associated point mutations can be corrected by these two BEs. And ABE, in particular, can correct up to 47% of the mutations [14, 15].

**Figure 1.**
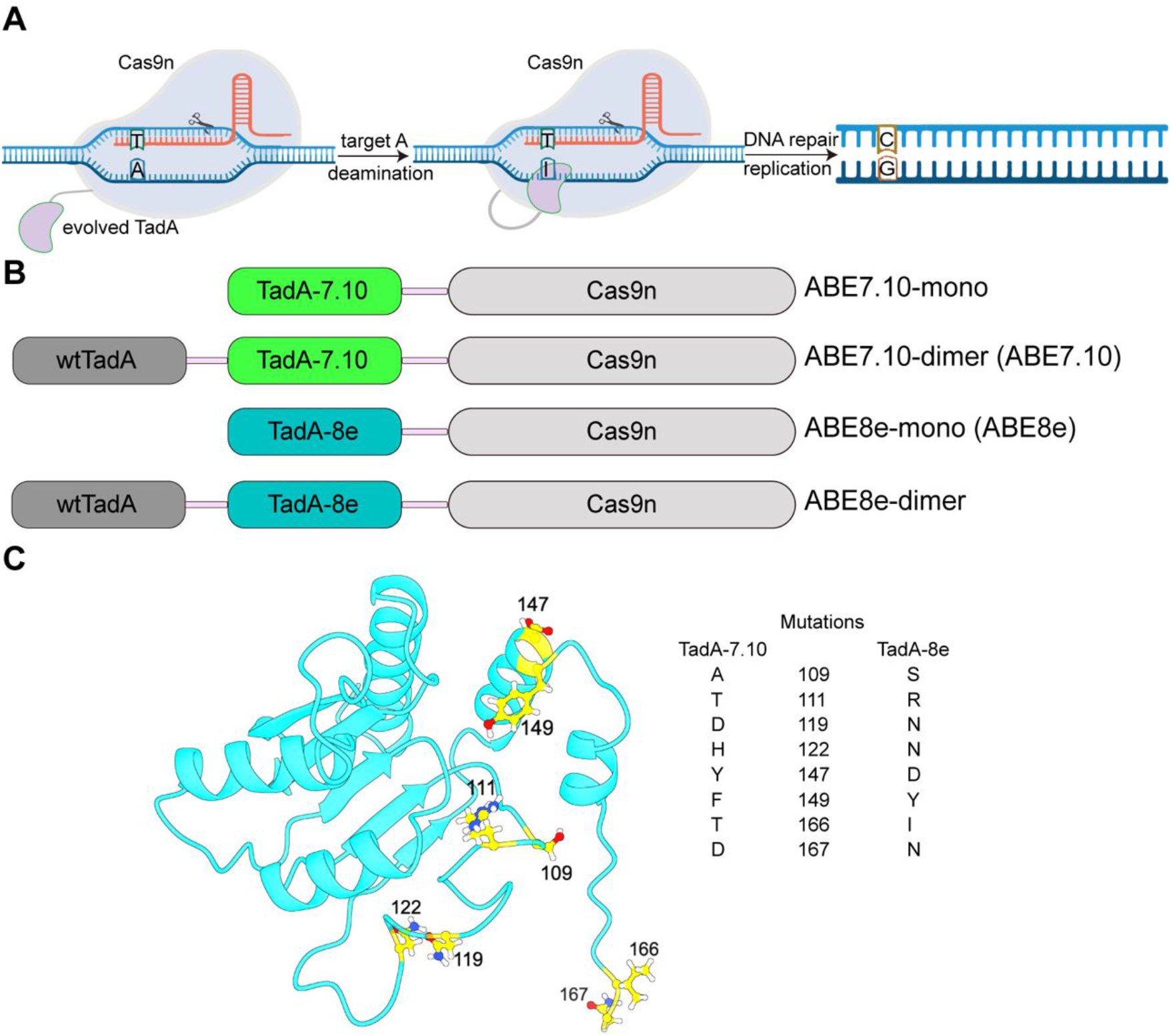
Overview of adenine base editing. (**A**) Schematic representation of ABE acting on DNA. Cas9 nickase fused to the evolved TadA. The programmed sgRNA binds to Cas to create a ssDNA R-loop, whereby the fused TadA converts exposed adenosine to inosine. The I:T is converted to G:C base pair after replication or additional DNA repair. **(B)** Domain architectures of four ABEs encoding wtTadA, evolved TadA-7.10 (green), or evolved TadA-8e (teal blue) N-terminally linked to Cas9n. **(C)** The eight directed-evolution mutations in TadA-8e that lead to the higher editing activity of ABE8e.

To develop ABEs, Liu and co-workers first used directed evolution to screen for variants of fusion proteins consisting of *E. coli* tRNA adenosine deaminase (TadA) and SpCas9 nickase (Cas9n), and successfully obtained an active BE called ABE7.10 [17]. ABE7.10 contains two TadAs: the wild-type *E. coli* TadA (wtTadA) and the evolved deaminase TadA-7.10 from wtTadA, and these two TadAs are fused to the SpCas9 nickase as wtTadA-TadA-7.10-Cas9n (Fig. 1B). It has been shown that in the absence of wtTadA, ABE7.10 editing activity is significantly reduced, indicating that two fused TadAs are required for this system [17, 22, 23]. However, the exact functional role of wtTadA remains unclear. In order to increase its activity and compatibility with other Cas effectors, additional rounds of directed evolution were carried out and then eight mutations were introduced into TadA-7.10 (Fig. 1C), resulting in a more active BE with a newly evolved TadA (TadA-8e), namely ABE8e [21]. Compared to ABE7.10, ABE8e requires only one deaminase (TadA-8e), but has the same editing activity in the absence of wtTadA, and its editing level at the same target sites was 3 to 11 times higher than that of ABE7.10. Thus, the eight directed-evolution mutations in TadA-8e significantly improve the editing activity, and also increase the compatibility of ABE8e with multiple Cas effectors [21].

As a rational basis, it is important to understand the mechanistic and functional roles of the directed-evolution mutations in ABEs in order to optimise them for efficient and precise editing by protein engineering. To this end, Rallapalli et al. have shown that the initial mutations in the evolved deaminase TadA can lead to intricate conformational changes in its structure [24]. Lapinate et al. have determined the first cryo-electron microscopy (cryo-EM) structure of ABE8e (PDB ID: 6VPC), and showed that the directed-evolution mutations could stabilise the DNA substrate in a constrained, transfer RNA-like conformation [22]. Recently, Rallapalli et al. also investigated the role of ABE mutations in RNA editing, and identified two important mutations affecting RNA-binding and catalytic activity [25]. Despite much effort on this subject, it remains unclear how the directed-evolution mutations in TadA-8e result in ABE8e having higher DNA editing activity than that of ABE7.10, and why ABE8e does not require wtTadA for its editing activity.

Here, we combine molecular dynamics (MD) simulations and experimental measurements to elucidate the molecular origin for the enhancement of DNA editing activity by the eight directed-evolution mutations in TadA-8e. MD simulations and microscale thermophoresis (MST) experiments showed that the DNA-binding affinity of TadA-8e is about twice that of TadA-7.10. We revealed that the electrostatic interactions are the main driving force for this, and the directed-evolution mutations increase the positive charge density in the DNA-binding region, thus increasing the electrostatic attractions of TadA-8e with the single-stranded DNA substrate. In addition, we found that the directed-evolution mutations also stabilise the thermal stability of TadA-8e, with a melting temperature (T_m_) higher than that of TadA-7.10 by about 12°C. Thus, these results highlight the functional roles of DNA-binding and protein stability in BEs.

## Results

### MD simulations of full-length ABEs

To understand how the directed-evolution mutations of ABE8e affect its base editing activity, it is necessary to investigate the 3D structure of its full-length model. To this end, we used the cryo-EM structure (PDB ID: 6VPC) as a template to build the full-length atomic model of ABE8e. However, in the cryo-EM structure the 32-aa peptide linker between Cas9n and TadA-8e and a certain number of NTS bases are missing. To complete these missing atoms, we first compared this structure with all Cas9-sgRNA-DNA ternary complexes available in the PDB, and found that the sgRNA-DNA conformation of 6VPC is similar to that in the active SpCas9 structures [26, 27] (Fig. 2A). Therefore, we used the NTS of the most similar structure (PDB ID: 5F9R) as a template to complete the missing NTS bases in ABE8e. We then constructed the 32-aa peptide linker using SWISS-MODEL (https://swissmodel.expasy.org), and thereby to obtain the full-length ABE8e model consisting of sgRNA, DNA, Cas9n and a single TadA-8e and the linker between them (Fig. 2B). This model contains all the missing amino-acids and bases in 6VPC and resembles a functional ABE8e in the cell. Finally, energy minimisation was performed to refine the built atoms in the model, and the refined model was named ABE8e-mono.

**Figure 2.**
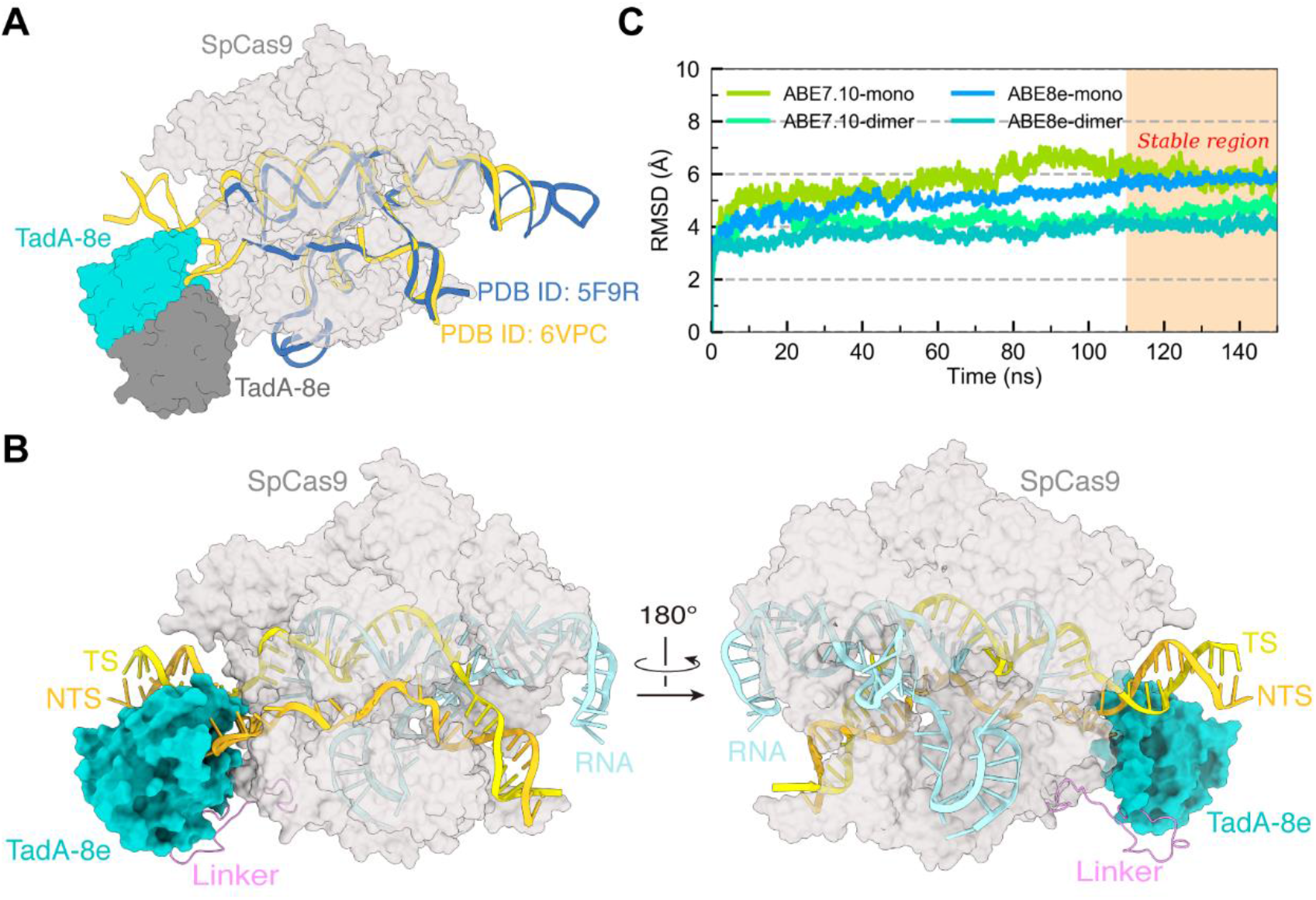
The full-length structure of ABE8e and RMSDs. **(A)** The sgRNA-DNA backbone alignment of ABE8e structure (PDB ID: 6VPC, orange) with SpCas9 structure (PDB ID: 5F9R, deep blue). **(B)** The full-length atomic model of ABE8e. SpCas9, light gray; RNA, cyan; TS, yellow; NTS, orange; TadA-8e, teal blue. **(C)** The time-dependent RMSDs of the ABEs.

As mentioned above, previous studies have shown that an additional wtTadA is needed for the ABE7.10 activity, but not for that of ABE8e [17, 21]. In order to understand the functional role of the wtTadA in ABE7.10, in addition to the above ABE8e-mono, we also constructed three other simulated models for detailed comparison (Fig. S1). The first is a fulllength ABE8e combined with an additional wtTadA (designated as ABE8e-dimer), and the second is ABE7.10 with only TadA-7.10 and without wtTadA (designated as ABE7.10-mono), and the third is ABE7.10 with TadA-7.10 and an additional wtTadA (designated as ABE7.10-dimer) (Fig. 1B).

To obtain equilibrated structures for the analyses, we performed all-atom MD simulations of about 150 ns for all four systems, as described in Materials and Methods. To assess whether the systems were equilibrated, we calculated the root mean square deviations (RMSDs) of the ABE backbone atoms using their initial structures as the references. As shown in Fig. 2C, the RMSDs of four systems displayed no significant changes after about 110-ns simulations, indicating that the simulated ABE complexes had been equilibrated. Therefore, the complex structures after 110 ns were used for the following analyses.

### TadA-8e has higher DNA-binding affinity

To reveal the differences in DNA binding between TadA-7.10 and TadA-8e, we analysed their equilibrated conformations of the TadAs and the DNA substrate. As expected, after 110-ns simulations TadA-8e in ABE8e-mono maintained in the substrate-bound state (Fig. 3A), and the distance from the active-site residue E59 to A26 of the substrate is ∼9.1 Å, slightly smaller than that in ABE8e-dimer (∼10.3 Å) (Fig. 3B). This implies that single TadA-8e in ABE8e is able to bind the DNA substrate. However, the corresponding distance of TadA-7.10 in ABE7.10-mono is ∼14.1 Å, significantly larger than that in ABE7.10-dimer (∼10.8 Å) and TadA-8e in ABE8e systems (Fig. 3B). Thus, the simulations suggest that the DNA-binding ability of TadA-7.10 in ABE7.10-mono is weak, so that an additional wtTadA is required to enhance this binding ability, such as in ABE7.10-dimer.

**Figure 3.**
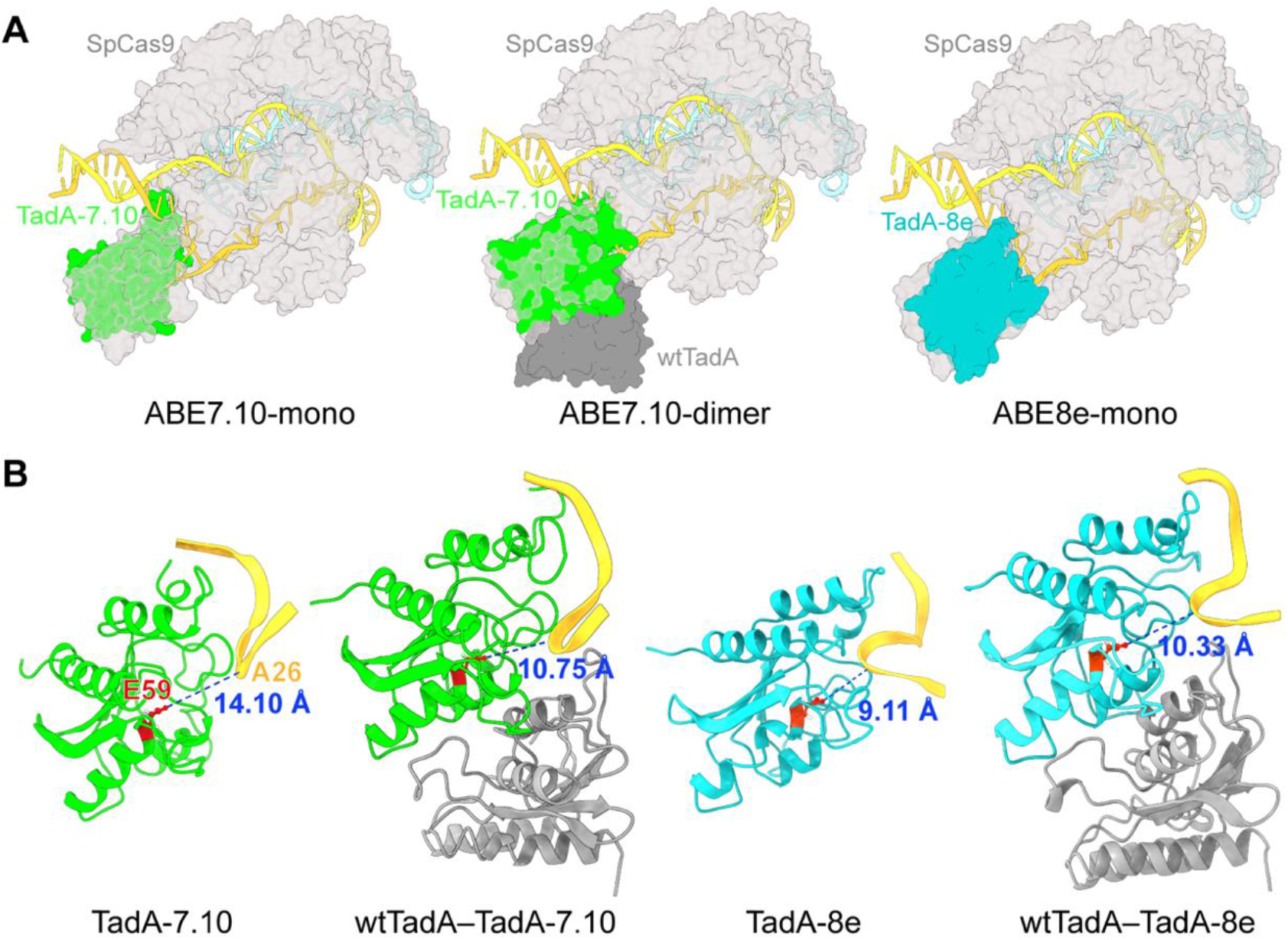
The conformational fluctuations of TadA-8e and TadA-7.10. **(A)** Alignment of the equilibrium and initial conformations of TadAs. The equilibrium conformations of TadA-7.10, wtTadA and TadA-8e are shown in green, grey, and teal blue, respectively. The initial conformations of TadAs are shown in light gray. **(B)** The distances from the active-site residue E59 of TadAs to the base A26 of the DNA substrate (blue dotted lines).

To verify the above hypothesis, we used the *g_mmpbsa* program [28] to calculate the binding energies for TadAs to the DNA substrate in the simulations. It is well known that the relative dielectric constant of the solute is a key parameter in the MM/PBSA calculation [28, 29]. Considering that the investigated ABEs are protein-DNA systems, to rationalise the comparisons, we used four different solute dielectric constants (2, 4, 6 and 8) for the calculations, the minimum 2 being commonly used for low polarised proteins [29, 30], and the maximum 8 being the DNA dielectric constant [31]. The calculated binding energies of TadAs to the ssDNA substrate (NTS) of the four simulated systems are listed in Table 1.

**Table 1.** Binding energies of TadA to NTS of DNA estimated using *g_mmpbsa* (in kcal/mol).

| Dielectric constant | TadA format | $\Delta E_{\text{elec}}$ | $\Delta E_{\text{vdW}}$ | $\Delta G_{\text{polar}}$ | $\Delta G_{\text{nonpolar}}$ | $\Delta G_{\text{binding}}$ |
| --- | --- | --- | --- | --- | --- | --- |
| $\epsilon = 2$ | TadA-7.10 | -777.84 | -91.34 | 292.50 | -10.64 | -587.42 |
|  | wtTadA–TadA-7.10 | -955.14 | -79.18 | 311.57 | -10.40 | -732.88 |
|  | TadA-8e | -1247.27 | -67.36 | 240.11 | -8.40 | -1082.99 |
|  | wtTadA–TadA-8e | -1434.52 | -71.59 | 438.25 | -9.38 | -1077.36 |
| $\epsilon = 4$ | TadA-7.10 | -388.84 | -91.31 | 281.53 | -10.64 | -209.30 |
|  | wtTadA–TadA-7.10 | -477.61 | -79.17 | 298.10 | -10.40 | -268.71 |
|  | TadA-8e | -623.53 | -67.33 | 236.89 | -8.40 | -462.58 |
|  | wtTadA–TadA-8e | -717.30 | -71.57 | 409.21 | -9.37 | -389.07 |
| $\epsilon = 6$ | TadA-7.10 | -259.32 | -91.34 | 272.37 | -10.64 | -88.86 |
|  | wtTadA–TadA-7.10 | -318.45 | -79.13 | 287.69 | -10.40 | -120.46 |
|  | TadA-8e | -415.74 | -67.35 | 234.75 | -8.40 | -256.74 |
|  | wtTadA–TadA-8e | -478.24 | -71.57 | 386.90 | -9.37 | -172.30 |
| $\epsilon = 8$ | TadA-7.10 | -194.42 | -91.40 | 264.70 | -10.63 | -31.65 |
|  | wtTadA–TadA-7.10 | -238.80 | -79.14 | 278.97 | -10.40 | -49.36 |
|  | TadA-8e | -311.82 | -67.34 | 233.38 | -8.40 | -154.10 |
|  | wtTadA–TadA-8e | -358.68 | -71.56 | 369.64 | -9.37 | -70.16 |

As shown in Table 1, in ABE7.10-mono and ABE8e-mono, the binding energy of TadA-8e with the ssDNA substrate was 2.0∼5.0 times that of TadA-7.10 at four different dielectric constants, indicating that TadA-8e has higher DNA-binding affinity than TadA-7.10. From the energies of ABE7.10-dimer, we found that wtTadA can assist TadA-7.10 with the binding to ssDNA; however, the total binding energies are still higher than those of single TadA-8e in ABE8e-mono, i.e., lower binding affinity. Interestingly, wtTadA in ABE8e-dimer does not contribute to the binding of TadA-8e to ssDNA, but instead introduces large polar solvent energy, resulting in the higher binding energy than that of TadA-8e, i.e., lower binding affinity. Consequently, the order of the binding affinities of TadAs to the ssDNA substrate is TadA-8e > wtTadA–TadA-8e > wtTadA–TadA-7.10 > TadA-7.10, which is in good agreement with the order of distances from their active sites to the substrates described above. Thus, wtTadA could increase the DNA-binding affinity of TadA-7.10 in ABE7.10, but does not contribute to that of TadA-8e; and, the directed-evolution mutations in TadA-8e significantly increase its binding capacity for the ssDNA substrate.

### Electrostatic attractions improve DNA-binding ability of TadA-8e

To determine the main forces driving the binding of TadAs to DNA, we analysed the components of the binding energies at all dielectric constants. As shown in Table 1 and Fig. S2, the binding between TadAs and the DNA substrate was largely driven by electrostatic attractions. Furthermore, the improved binding energy in TadA-8e was also largely due to the enhanced electrostatic attractions. To uncover the structural basis for the difference in electrostatic interactions between TadA-8e and TadA-7.10, we calculated their surface electrostatic potentials using the APBS program [32].

As showed in Fig. 4, the DNA-binding regions of the TadAs are distributed with many positively charged residues; however, the surface potentials indicate that the positive charge density and area on the surface of TadA-8e are higher and larger than those in TadA-7.10. It is therefore reasonable to infer that when a negatively charged ssDNA substrate approaches, the stronger electrostatic attractions of TadA-8e could capture the substrate more efficiently, and this could explain the higher deamination rate of TadA-8e [21, 22]. In addition, there are many positively charged residues distributed in the interface region of wtTadA with the DNA backbone (Fig. 4), which also suggests that wtTadA has the potential to assist TadA-7.10 in binding to the substrate, as indicated by the results in Table 1.

**Figure 4.**
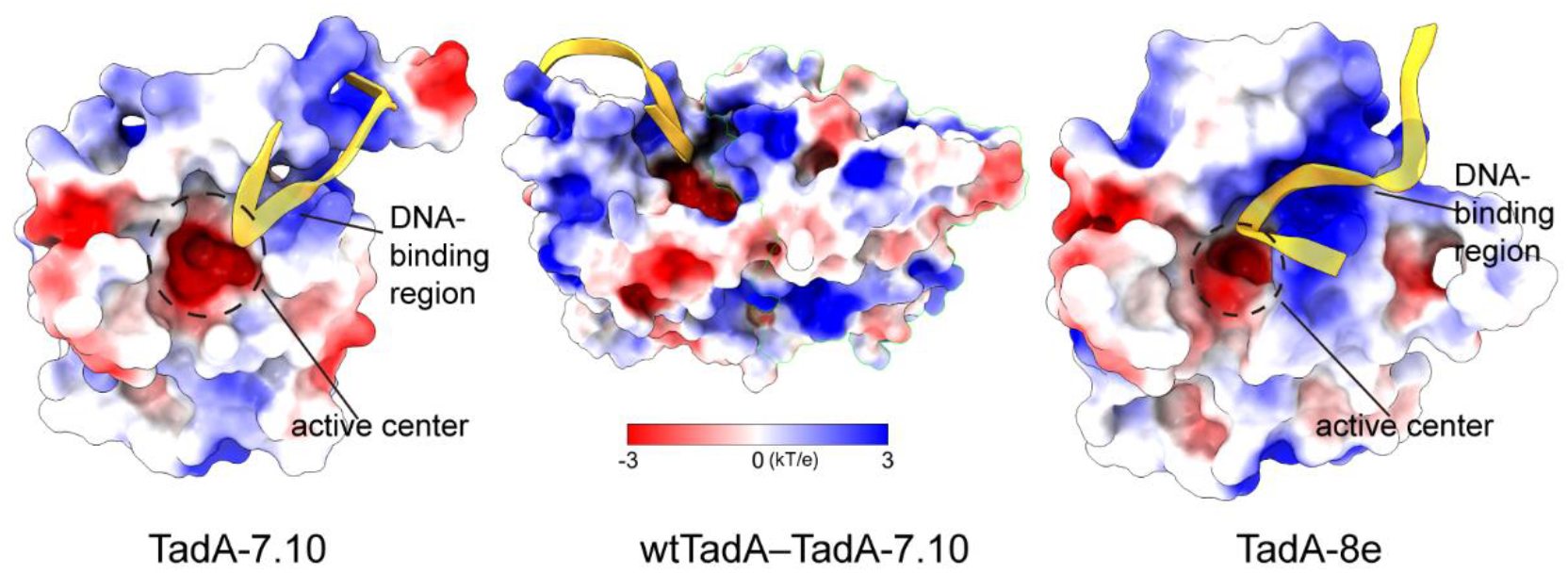
The electrostatic potentials on the surfaces of TadA-7.10, wtTadA-TadA-7.10 and TadA-8e. The electrostatic potentials were calculated using the PDB2PQR server and PyMOL with the *APBS* plugin on a scale from -3 to 3 kT/e.

Meanwhile, in TadA-7.10, the negatively charged area of the active site is over-exposed, which could repel the negative ssDNA and thus prevent the ssDNA substrate from accessing the active site. In contrast, in TadA-8e, the ssDNA is mainly bound to the positive potential surface and its conformation is strongly restricted (Fig. 4). This is certainly favourable for the active site to effectively catalyse the editing reactions. Consequently, compared to that of TadA-7.10, the electrostatic potentials at the DNA binding region of TadA-8e favour stronger binding to ssDNA, thus increasing the DNA binding affinity of ABE8e.

### Mutations increase positive charge density in the DNA-binding region of TadA-8e

Since TadA-8e was obtained by mutating eight residues of TadA-7.10, the higher DNA-binding affinity of TadA-8e should be determined by these mutations. To quantitatively characterise the contributions of the mutations to the DNA binding, we calculated the binding energy difference (ΔΔG) for each amino acid between TadA-8e and TadA-7.10. As mentioned above, TadAs are proteins with highly polar residues on the surface, so the following energy analysis takes the dielectric constant of 6 as an example for the calculations, and the results are shown in Fig. 5A. As can be seen, the mutations with the significant contributions (ΔΔG < -20 kcal/mol) are

**Figure 5.**
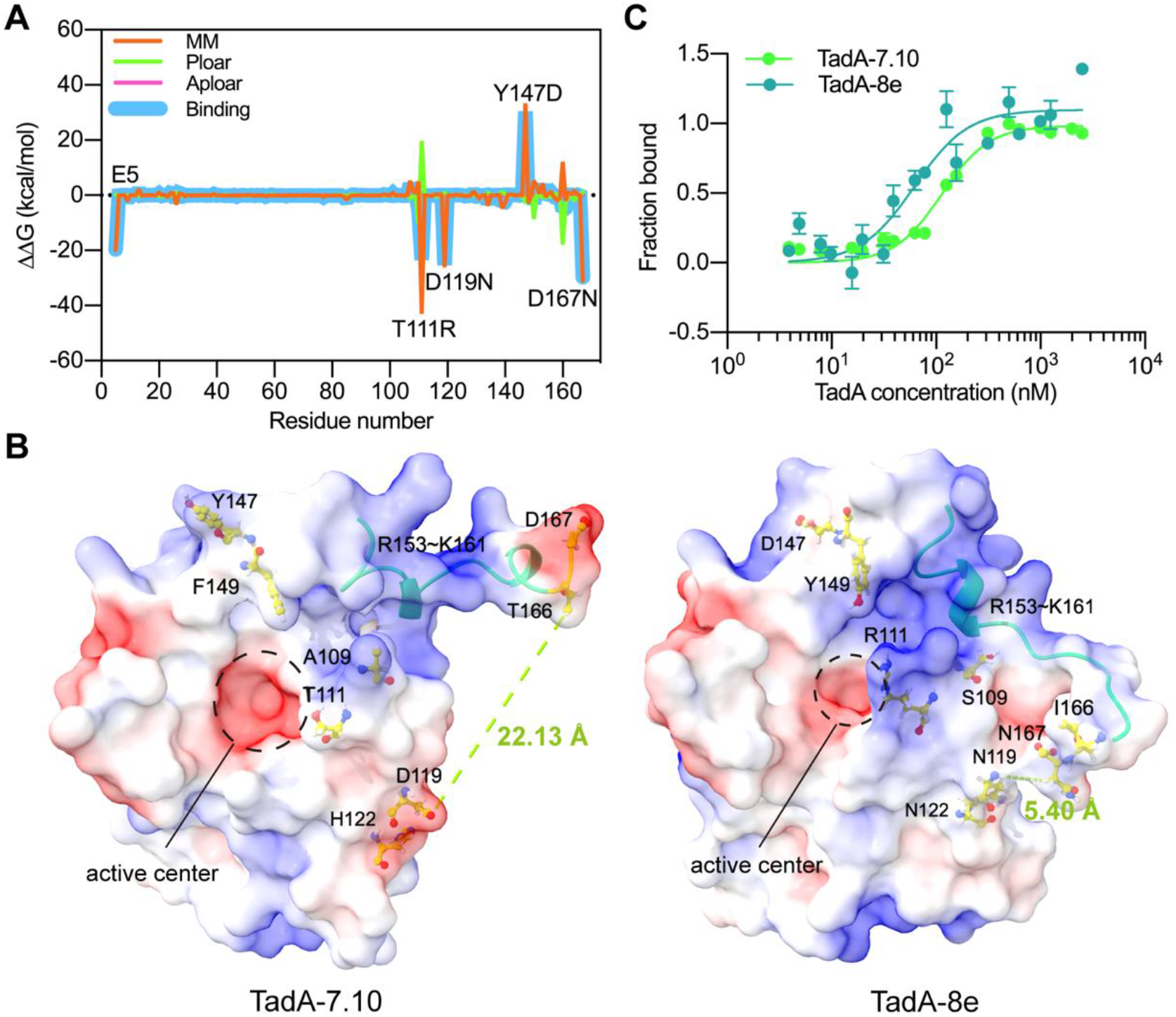
The electrostatic contribution of mutations to TadA-8e binding DNA and MST experiments. **(A)** The ΔΔG value per residue between TadA-8e and TadA-7.10. **(B)** The electrostatic potential patches of mutations. Coulombic potentials (red, negative; blue, positive) were calculated with the APBS program. **(C)** The *K*_d_ values of TadA-7.10 and TadA-8e are 113.40 ± 0. 57 nM and 62.10 ± 0. 13 nM, respectively.

T111R, D119N and D167N (Table S1), and the main energy component is electrostatic interactions. Among them, N119 and N167 eliminate the electrostatic repulsions and the positively charged residue R111 directly increases the electrostatic attractions. Indeed, R111 is the amino acid that contributes most to the binding (Table S1), which explains why the editing activity of ABE8e was dramatically reduced after R111 was backmutated to the original T [22]. To reveal the effects of the mutations on the surface electrostatic potentials of TadA-8e, we analysed the electrostatic distributions caused by these mutations. As shown in Fig. 5B, in TadA-8e, R111 increases the positively charged area in the DNA-binding region, again indicating that R111 is a key residue for the DNA-binding affinity. In addition, in TadA-7.10, a possible electrostatic repulsion between D119 and D167 causes the peptide R153∼K161 to move away from the active site, resulting in a disruption of the continuous positive potential surface (Fig. 5B). Thus, if D119 and D167 were mutated to N, the repulsion effect could be eliminated, then N167 would approach N119, and the R153∼K161 peptide and R111 could form a continuous surface of positive potentials, expanding the area of positive charges in the DNA-binding region. Most likely, R111, N119 and N167 enhance the electrostatic attractions between TadA-8e and ssDNA by expanding the positively charged area, thus further improving the DNA-binding affinity.

To verify that TadA-8e has a higher DNA-binding affinity, we next performed MST measurements to experimentally determine the binding affinities of TadA-7.10 and TadA-8e to the ssDNA substrate. First, we expressed and purified the inactive TadA-7.10 and TadA-8e proteins (Fig. S3). And the ssDNA substrate was derived from the NTS in the structure 6VPC. Since TadAs only act on ssDNA, to test whether the NTS used is single-stranded, we predicted its secondary structure using the RNAStructure website (http://rna.urmc.rochester.edu/RNAstructureWeb/) [33], and found that it is easy to form base-paired structure (Fig. S4A). Therefore, we mutated five bases each at its two ends and obtained a 19-nt DNA substrate (TTCTCTTCC<u>A</u>CTTTCTTTT) as the ssDNA substrate in the MST measurements. The MST results showed that the equilibrium dissociation constants (*K*_d_) of TadA-7.10 and TadA-8e were 113.40 ± 0.57 nM and 62.10 ± 0.13 nM, respectively (Fig. 5C). Therefore, in agreement with the calculations, our MST experiments also showed that the DNA-binding affinity of TadA-8e is approximately twice that of TadA-7.10, strongly supporting that the directed-evolution mutations of TadA-8e improve its DNA-binding affinity.

### Directed-evolution mutations also improve protein stability of TadA-8e

As mentioned above, electrostatics is the main driving force for TadAs-DNA binding. Electrostatics not only affect the interactions between two molecules, but also have influence on the physicochemical properties of a protein, such as the thermal stability [34-36]. To investigate the effects of electrostatics on the stability of TadA-7.10 and TadA-8e, we compared their electrostatic potentials on the protein surfaces (Movies S1 and S2). As seen, the potential surface of TadA-7.10 was relatively loose, while that of TadA-8e is smooth, suggesting that the structure of TadA-8e is more compact, and its electrostatic potentials on surface appears to be more optimal for its editing function. To quantify the effects, we used the TKSA-MC server (http://tksamc.df.ibilce.unesp.br/) to calculate the electrostatic contributions of polar-charged residues to the folding free energies of TadA-7.10 and TadA-8e, respectively [37]. The results showed that the total free-energy contribution to TadA-8e (-24.31 kcal/mol) is lower than that to TadA-7.10 (-17.29 kcal/mol) (Fig. 6A), suggesting that TadA-8e is more stable than TadA-7.10 due to the directed mutations.

**Figure 6.**
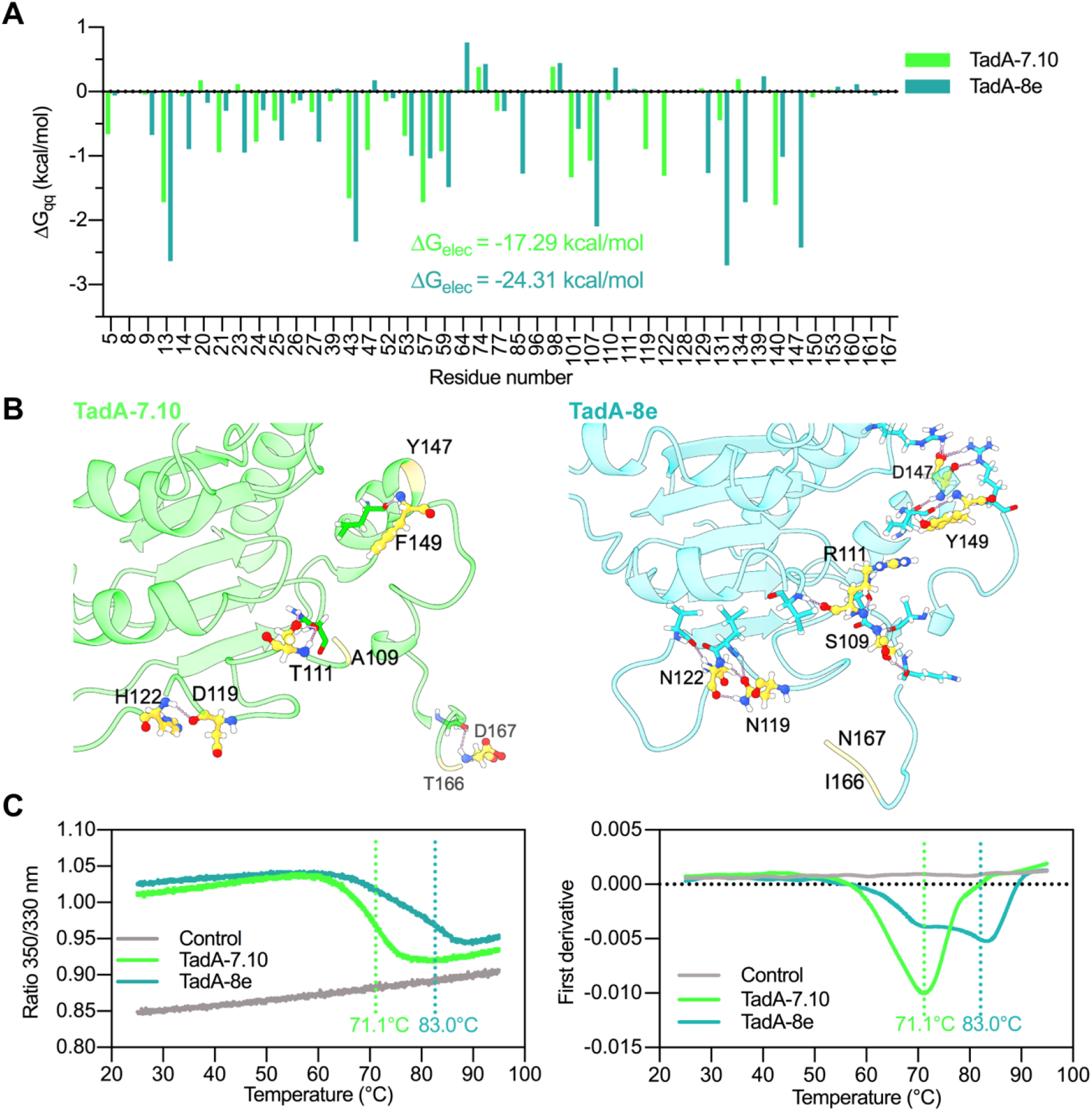
The effect of mutations on the stability of TadA-7.10 and TadA-8e. **(A)** The charge-charge interaction energy (ΔGqq) calculated by the TKSA-MC model for each ionizable residue of TadA-7.10 and TadA-8e in pH 7.0, 310 K. **(B)** Hydrogen bond distribution of mutations in TadA-7.10 and TadA-8e, respectively. The mutations in TadA-7.10 and TadA-8e are represented in yellow ball stick. Hydrogen bonds are shown as pink dotted lines. **(C)** Nano-differential scanning fluorimetry (nanoDSF) curves (left) and their first derivative (right) of TadA-7.10 (green) and TadA-8e (teal blue).

In addition to the electrostatic optimization, hydrogen bonding also contributes to protein stability [38, 39]. Therefore, we further analysed the hydrogen bonding patterns in two proteins. As shown in Fig. 6B, mutations in TadA-8e form more hydrogen bonds than those in TasA-7.10. In fact, corresponding bonding sites in TadA-7.10 form only a few hydrogen bonds (Table S2). Again, this suggests that the directed-evolution mutations of TadA-8e allow for more hydrogen bonding interactions, further stabilizing the protein structure.

To verify the above computational analyses, we next performed thermal shift assays using nanoscale differential scanning fluorimetry (nanoDSF)[40] to measure the melting temperature (T_m_) of two proteins. NanoDSF can characterise protein stability, folding, and aggregation of proteins by monitoring small changes in intrinsic tryptophan fluorescence (330 nm, 350 nm) during protein unfolding. As shown in Fig. 6C, T_m_ of TadA-8e is ∼ 83.0°C, while that of TadA-7.10 is about 71.1°C. Thus, the thermal stability of TadA-8e is enhanced by ∼12°C (T_m_), providing strong evidence that TadA-8e has higher thermostability than TadA-7.10. In conclusion, the eight directed-evolution mutations in TadA-8e improve its protein stability by optimising protein electrostatics and forming more hydrogen bonds.

## Discussion

In the present study, we have combined MD simulations and experimental measurements to understand how the eight directed-evolution mutations in the TadA-8e deaminase of ABE8e enhance its base editing activity. We showed that TadA-8e has a higher DNA-binding affinity than the original TadA-7.10 without these directed-evolution mutations, and that electrostatic interactions are the main driving force for TadA-8e binding to ssDNA. We also showed that the directed-evolution mutations increase the positive charge density in the DNA-binding region of TadA-8e, thereby enhancing the electrostatic attraction with the NTS of DNA. Moreover, we found that the directed-evolution mutations significantly improved the thermal stability of TadA-8e. Taken together, our data elucidate the molecular origin of the activity enhancement by the directed-evolution mutations of ABE8e, and demonstrate that the base editing activities of ABEs are closely related to the DNA-binding affinity and protein stability of their deaminases.

In our study, we have shown that due to the enhanced DNA-binding induced by the directed-evolution mutations, only one deaminase (TadA-8e) is sufficient for ABE8e to attract the NTS of DNA to its active site. In contrast, for ABE-7.10, an additional deaminase (wtTadA) is required to support such binding in TadA-7.10. This explains the experimental observation that wtTadA is required for ABE7.10 to carry out its base editing activity [17, 22, 23]. Of course, it remains difficult to quantify the contributions of the enhanced DNA-binding affinity to the base editing activity; many studies have shown a positive correlation between the substrate-binding affinity and the nuclease activity [41-46]. For example, the DNA-binding affinity and specificity of the zinc finger nucleases are the main determinants of their effective activity in human cells [42]. Similarly, the increased binding affinity of *Lb*Cas12a for its hairpin-forming substrates results in increased *trans*-cleavage [43]; and the high affinity of activation-induced deaminase (AID) for DNA substrates is required for its efficient deamination [45]. In agreement with our study, Rallapalli et al. also showed that the D108N mutation of TadA* improves its early editing kinetics by increasing the substrate binding [24]. As we have pointed out, this also explains why the back mutation of the key DNA-binding residue R111 to the origin T dramatically reduced the editing activity of ABE8e. All these results suggest that the high affinity of TadA-8e for the target DNA is a major determinant of its base editing activity. Of course, further studies are needed to clarify the relationship between the DNA-binding affinity and the base editing activity.

Many studies have shown that enzymes with higher thermal stability tend to be more active than those with lower thermal stability at the same temperature [47]. One possible explanation is that for more stable enzymes, more molecules could survive the potential thermal unfolding at the given temperature and then have the active, functional structure for the activity. Here, we have shown an unexpected finding that the protein stability of TadA-8e is increased simultaneously with its editing activity. Thus, our study provides new evidence for that directed evolution can effectively improve protein activity and stability [48-50]. This may provide useful guidance for optimising enzymes with high activity and stability by learning the mutation patterns of directed-evolution variants.

In summary, ABEs are important molecular machines for precise genome editing. In the past, directed evolution has been a major approach to their optimisation. Therefore, it is very important to elucidate the molecular origin of the directed-evolution mutations in order to improve the ABE activities. Here, we combined MD simulations and experimental measurements to reveal how the directed-evolution mutations in TadA-8e improve the base editing activity of ABE8e. Our results indicate that the base editing activities of ABEs are closely related to their DNA-binding affinity and protein stability. Therefore, our study provides not only new mechanistic insights into the molecular machines of base editing, but also useful guidance for their rational optimisation for efficient and precise genome editing.

## Materials and Methods

### Structure construction

The cryo-EM structure of ABE8e (PDB ID: 6VPC) was used to construct all the full-length models of ABEs for the MD simulations. The missing atoms and loops were constructed using homology modelling at SWISS-MODEL (https://swissmodel.expasy.org). The TadA-7.10 structure was generated by introducing point mutations into the TadA-8e structure using the *mutagenesis* plugin available in PyMOL. For the construction of the wtTadA structure, the *E. coli* TadA deaminase (PDB ID: 1Z3A) was used as a modelling template [51]. The surface electrostatic potentials for TadA-8e and TadA-7.10 were calculated using the *APBS* plugin available in PyMOL [32, 52]. All visualisations and interaction analyses were performed using ChimeraX [53, 54] and GROMACS software [55, 56].

### MD simulations

All MD simulations were carried out using GROMACS. The topology and coordinate files were generated using the *pdb2gmx* program with parameters from the AMBER99SB-ILDN force field [57]. The complex was placed in the center of a cubic box with an edge length of ∼15 Å from the protein surface to the box boundary, and then solvated with TIP3P water molecules [58]. Specific numbers of Na^+^ and Cl^−^ ions were added to the system to neutralise the complex charge and the ion concentration was set to 0.15 M. To optimise the system, energy minimisation was performed using the steepest descent algorithm for a maximum of 2500 steps or until the maximum force < 1000 kJ·mol ^−1^·nm ^−1^. The system was then heated to 310 K for 100 ps with a time step of 2 fs by constant NVT equilibration. Subsequently, constant NPT simulations of 500 ps were performed to equilibrate the system at a pressure of 1.0 bar. The simulated temperature and pressure were maintained at 310 K and 1.0 bar using the V-rescale temperature and Parrinello–Rahman pressure coupling methods, respectively [59, 60]. During the simulations, all bonds were constrained using the LINCS method [61], and the periodic boundary conditions (PBC) were applied in all three dimensions. The short-range non-bonded interactions were calculated for the atom pairs within a cut-off of 14 Å, while the long-range electrostatic interactions were calculated using the Particle Mesh Ewald (PME) method [62]. Finally, production simulations of ∼150 ns were performed for the following analyses, so that the coordinates of the system atoms were recorded per 100 ps.

### Calculation of binding energies

The *g_mmpbsa* package was used to calculate binding energy between the TadAs and NTS by the molecular mechanics Poisson–Boltzmann surface area (MM–PBSA) method [28, 63]. The energy components E_MM_, G_polar_, and G_nonpolar_ of each complex were calculated for the snapshots extracted from the MD production simulations from 110 to 150 ns (41 snapshots). The electrostatic energy, van der Waals energy and polar solvation energy were calculated using the Poisson–Boltzmann solver (APBS) method, and the non-polar energy was calculated by the solvent-accessible surface area (SASA) method [64]. The dielectric constant of the solvent was set to 80, and four values (2, 4, 6 and 8) were used for biomolecules [29]. The Python script *MmPbSaStat*.*py* was used for the MM-PBSA calculation and *MmPbSaDecomp*.*py* was used to calculate the contributions of individual residues in the TadAs.

### Protein expression and purification

TadA-7.10 and TadA-8e, bearing an N-terminal non-cleavable His_6_-tag, were expressed in *E. coli* BL21 (DE3). Cells were grown in LB containing 50 *μ*g/mL kanamycin and 25 *μ*g/mL chloramphenicol at 37°C to an OD_600_ of approximately 1, and were induced with 0.5 mM IPTG. Growth was continued at 25°C for 12 h. Cells were harvested by centrifugation and resuspended in buffer: 20 mM HEPES pH7.5, 500 mM KCl, 20 mM imidazole, 10% glycerol, 2 mM β-mercaptoethanol (β-ME) supplemented with protease inhibitor 1 mM PMSF, and lysed by sonication on ice. After centrifugation at 8,000 rpm for 1 h, the supernatant was loaded onto Poly-Prep chromatography columns (BIO-RAD) containing Ni-NTA agarose (QIAGEN), and proteins were eluted with a gradient of buffer supplemented with 500 mM imidazole. The eluted proteins were further purified by size exclusion chromatography using a Superdex 200 Increase 10/300 GL column developed in 20 mM HEPES pH7.5, 500 mM KCl buffer. Final purity was verified by the SDS-PAGE analysis. Eluted proteins were flash-frozen in liquid nitrogen and stored at −80 °C.

### Microscale thermophoresis (MST)

MST experiments to measure the binding affinities between TadAs and the 19-nt DNA substrate were performed on a Monolith NT.115 system (NanoTemper Technologies, Germany) using the nano BLUE detector. The MST experiments were performed using 15% LED-power and 40% IR-laser power. Laser on and off times were set to 30 s and 5 s, respectively. The 19-nt DNA labelled with 6-FAM at the 5’-end was purchased from Sangon Biotech Co., Ltd. (Shanghai, China). A final concentration of 100 nM of 19-nt DNA was used and TadAs were diluted 1:1 starting at 4000 nM and 5000 nM in the binding buffer: 20 mM HEPES pH 7.5, 150 Mm KCl, 10 mM MgCl_2_ 1mM DTT, supplemented with 0.05% Tween-20. Samples were incubated at room temperature for 120 min and subsequently filled into capillaries (NanoTemper Technologies) for the measurements. The error between the fluorescence intensities of all capillaries is no more than 10%, i.e.,1200-1320 (Fig. S4B), and data points outside this range were excluded. Measurements were performed at least three times, and the resulting data were analysed using the MO.Affinity analysis software (NanoTemper Technologies).

### Thermal shift assay

The melting temperature T_m_ of a TadA protein was determined using nano differential scanning fluorimetry (nanoDSF,Nanotemper Prometheus NT.48 system) [40]. The temperature was increased from 25 to 95°C at a ramp rate of 1°C/min. The excitation wavelength was set at 280 nm and the instrument monitored the emission fluorescence at 350 nm and 330 nm. In the measurements, the recorded ratio of the emission intensities (Em_350nm_/Em_330nm_) represents the change in tryptophan fluorescence. The T_m_ value for each experiment of the given TadA was then automatically calculated using the PR.ThermControl software by plotting the ratiometric measurement of the fluorescence signal against the increasing temperature.

## Supporting Information

*The online version contains supplementary material available at:*

Supporting figures and tables

Movie S1. The electrostatic potential on TadA-7.10 surface

Movie S2. The electrostatic potential on TadA-8e surface

## Acknowledgements

We thank the Shanghai Supercomputer Center for providing computational resources.

## Authors’ contributions

Conceptualization: H.Z. and Q.H. Methodology: H.Z., L.W., and M.S. Investigation: H.Z., X.J., and Q.Q. Visualization: H.Z. and Q. H. Supervision: Q.H. Writing (original draft): H.Z. and Q.H. Writing (review and editing): H.Z. and Q.H.

## Availability of data and materials

All data needed to evaluate the conclusions in the paper are present in the paper and/or the Supplementary information. Additional data related to this paper may be requested from the authors.

